# Glomerulosclerosis and kidney failure in a mouse model of monoclonal immunoglobulin light-chain deposition disease

**DOI:** 10.1101/624650

**Authors:** Sébastien Bender, Maria Victoria Ayala, Amélie Bonaud, Vincent Javaugue, Claire Carrion, Christelle Oblet, Alexia Rinsant, Nathalie Quellard, Sihem Kaaki, Zeliha Oruc, François Boyer, Agnès Paquet, Nicolas Pons, Bastien Hervé, Mohamad Omar Ashi, Arnaud Jaccard, Laurent Delpy, Guy Touchard, Michel Cogné, Frank Bridoux, Christophe Sirac

## Abstract

Light chain deposition disease (LCDD) is a rare disorder characterized by glomerular and peritubular amorphous deposits of a monoclonal immunoglobulin (Ig) light chain (LC), leading to nodular glomerulosclerosis and nephrotic syndrome. We developed a transgenic model using site-directed insertion of the variable domain of a pathogenic human LC gene into the mouse Ig kappa locus, ensuring its production by all plasma cells. High free LC levels were achieved after backcrossing with mice presenting increased plasma cell differentiation and no Ig heavy chain (HC) production. Our mouse model recapitulates the characteristic features of LCDD, including progressive glomerulosclerosis, nephrotic-range proteinuria and finally, kidney failure. The variable domain of the LC bears alone the structural properties involved in its pathogenicity. RNA sequencing conducted on plasma cells demonstrated that LCDD LC induces endoplasmic reticulum stress, likely accounting for the high efficiency of proteasome inhibitor-based therapy. Accordingly, reduction of circulating pathogenic LC was efficiently achieved and not only preserved renal function, but partially reversed kidney lesions. Finally, transcriptome analysis of pre-sclerotic glomeruli revealed that proliferation and extracellular matrix remodelling represented the first steps of glomerulosclerosis, paving the way for future therapeutic strategies in LCDD and other kidney diseases featuring diffuse glomerulosclerosis, particularly diabetic nephropathy.

## Introduction

Monoclonal gammopathies of renal significance (MGRS) are characterized by renal lesions due to monoclonal immunoglobulins produced by a non-malignant B or plasma cell clone (1). Depending on the type and structural properties of the Ig, they comprise tubulopathies caused by LC accumulating in the proximal or distal tubules, or glomerulopathies, most frequently AL amyloidosis and Randall-type monoclonal Ig deposition disease (MIDD) (2). MIDD are characterized by linear amorphous deposits of a monoclonal LC (LCDD) or a truncated HC (HCDD) along the tubular and glomerular basement membranes (BMs), around arteriolar myocytes and in the mesangium, eventually leading to diabetic-like nodular glomerulosclerosis. Clinical manifestations include glomerular proteinuria with progressive kidney failure (3–5). In LCDD, the most frequent form of MIDD, involved LCs are mostly of the kappa isotype, with a striking overrepresentation of the Vk4 variable domain characterized by a long CDR1. Other peculiarities of LCDD LCs are due to somatic hypermutations and include polar to hydrophobic amino-acids replacements, small truncations or abnormal glycosylations (3, 4, 6–10). Moreover, LCDD LCs are characterized by the constant high isoelectric point (pI) of their V domain compared to LCs involved in AL amyloidosis (11). This feature was also confirmed in HCDD and could account for the high avidity of positively charged LCs for anionic heparan sulfates of basement membranes (5). However, little is known about the pathogenic mechanisms involved in glomerular damage leading to proteinuria and glomerulosclerosis. Herrera and colleagues have shown, first *in vitro* then *in vivo*, the acquisition of a myofibroblast phenotype by mesangial cells exposed to purified LC from patients with LCDD (12–15). These changes came along with the increased production of PDGFβ and TGFβ, resulting in excessive generation of extracellular matrix (ECM) proteins. Similarly to diabetic nephropathy (16), ECM remodelling is supposed to contribute to the glomerular alterations observed in MIDD. However, since most of these experiments were performed *in vitro* and relied on acute exposure to pathogenic LC, they deserve to be confirmed in a model reproducing the progressive development of glomerular lesions observed in MIDD. Accordingly, we recently developed a transgenic model of MIDD in which the truncated HC gene from a HCDD patient was introduced into the mouse Ig kappa locus to obtain sustained and continuous production of the pathogenic human Ig by all plasma cells (17). This model reproduced the main pathologic features of HCDD, including amorphous linear deposits along tubular and glomerular basements membranes, but neither glomerulosclerosis nor kidney dysfunction was observed, likely due to the very low level of circulating pathogenic human HC in mice. In the present study, we developed a LCDD mouse model, κF-DH, using targeted insertion of a human LC gene into the mouse kappa locus coupled to a deletion of the HC genes to obtain high levels of circulating pathogenic free LC (FLC) (18). This strategy proved efficient with the induction of a full-blown LCDD and gave new insights into the pathophysiology of the disease and into the toxicity of monoclonal LCs for plasma cells.

## Results

### An efficient transgenic strategy to produce high amounts of human pathogenic LCs in mouse

To obtain a high production of the pathogenic human kappa LC, we chose to use a Ig kappa knock-in strategy that proved successful in previous models of Fanconi syndrome and HCDD (17, 19). We introduced the VJ exon sequence obtained from a patient (κF) with biopsy-proven LCDD in place of the Jκ segments, precluding all further mouse kappa rearrangements in homozygous mice (Figure 1A). This approach allows an efficient and physiological production of the LC by all B and plasma cells, mimicking the monoclonal status of the patient. Transgenic LCs are chimeric proteins composed of the human variable domain associated to the endogenous mouse constant region and were shown to associate normally to mouse heavy chains (19, 20). Consequently, in the aim to increase the production of FLC, we further crossed these mice with the DH-LMP2A mice (“DH mice” hereafter) in which the HC locus was invalidated by the targeted insertion of the Epstein Barr virus protein LMP2A that mimics BCR signalling and allows complete B cell development (Figure 1B)(21). DH mice feature increased plasma cell differentiation despite the absence of endogenous HCs (22). Consequently, LCDD mice, corresponding to the double homozygous DH and κF strain (κF-DH), closely recapitulate the features of monoclonal gammopathy with an elevated number of plasma cells producing the human/mouse chimeric FLC (Figure 1, C and D). Using this strategy, we obtained serum FLC levels similar to those observed in patients with LCDD (23). Serum production of FLC progressively increased with age and lasted during the life-span of the animals. Bence-Jones proteinuria was readily detectable in 5 month-old mice ((Figure 1E).

**Figure 1:**
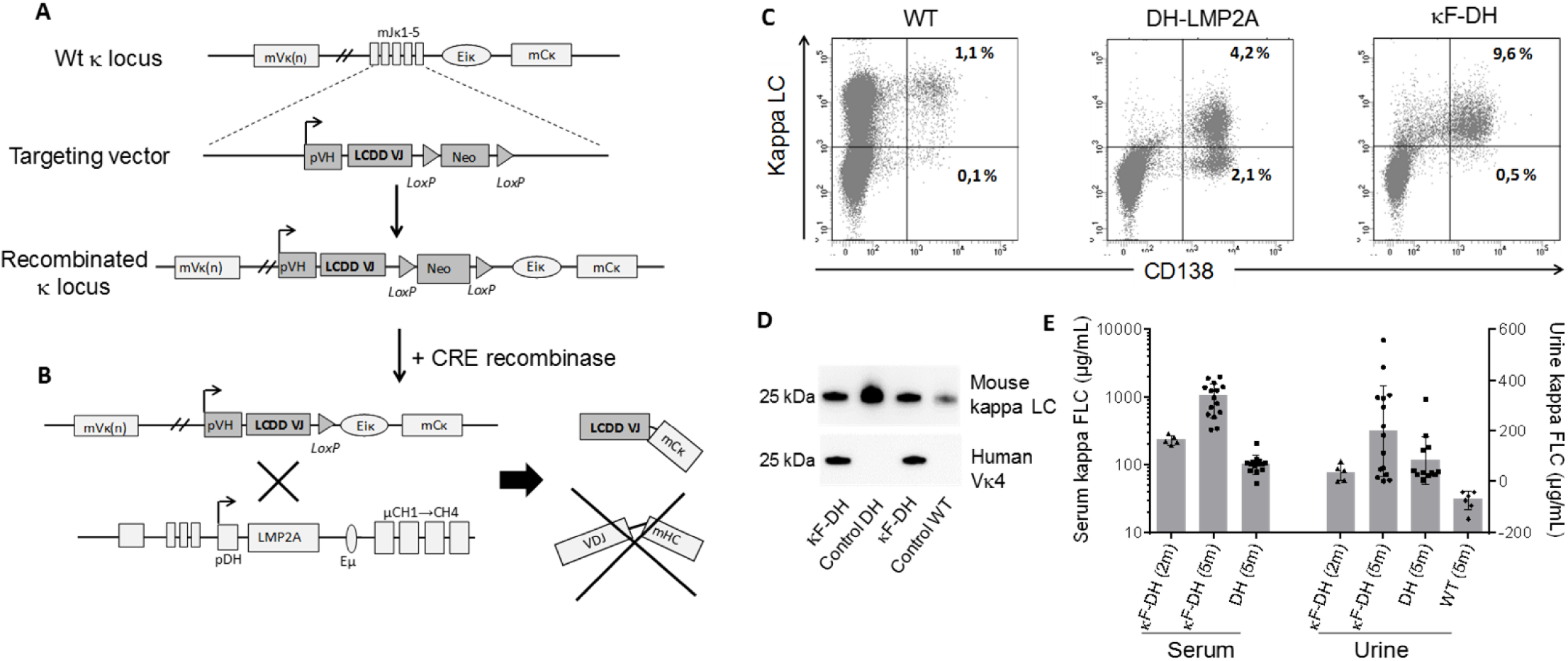
Strategy to produce high amounts of a human kappa light chain in mouse. (A) The strategy consists in replacing the Jk segments in a WT unarranged kappa locus by a human VJ exon/neomycin cassette. The absence of the Jk segments in the newly recombined kappa locus blocks all possibilities of endogenous rearrangements. A cre-mediated deletion of the neomycin resistance gene leads to the production of a chimeric human VJ/mouse kappa constant light chain. (B) Breeding with DH-LMP2A (DH) mice enables the production of only free light chain by B and plasma cells. (C) Flow cytometry analysis of plasma cells isolated from spleen and stained with anti-mouse CD138 and anti-mouse kappa antibodies. Note the increase of plasma cells in DH models compared to WT and that nearly all plasma cells are kappa+/CD138+ in κF-DH mice. (D) Western blot analysis of the produced kappa light chains in sera of κF-DH, DH and WT mouse with an anti-mouse kappa antibody (top) and with an anti-human Vκ4 variable domain (bottom). The bands appears at the expected size of 25 kDa and the anti-human Vκ4 antibody reveals only the chimeric kappa light chain of the κF-DH mice. (E) Serum and urine levels (in µg/mL) of free kappa light chains in 2 months and 5 months-old κF-DH mice compared to 5 months-old DH and WT mice. The free kappa light chain levels increase in κF-DH mice with age. Serum results are expressed in log scale, means are ± SEM.

### Severe kidney failure induces premature death of LCDD mice

Two cohorts of κF-DH mice (total of 12 mice composed of 7 males and 5 females) were monitored daily from the fifth month for external signs of morbidity (see methods). No signs of morbidity were observed in mice before 6 months of age. The first mouse was sacrificed at 6 months due to visible alteration of general condition. Then the entire cohorts had to be sacrificed within the first 14 months, with a median survival of 8.5 months (Figure 2A). DH mice were used as control, since, similarly to κF-DH mice, they do not produce complete antibodies (10 mice, 5males and 5 females). At 14 months, in our pathogen-free facility, all DH mice were healthy (Figure 2A). Having demonstrated that the early death of κF-DH mice was not due to immunodeficiency, we analysed their renal function. We first explored the progression of albuminuria with age (Figure 2B). No albuminuria was observed in 2 or 6 month-old κF-DH mice as compared to control mice. At 8 months of age, mice without signs of morbidity presented a significant increase in albuminuria compared to controls (Figure 2B). Mice sacrificed at humane endpoints (median survival: 8.5 months) showed heavy albuminuria, reaching up to 24.80 g/L (mean 8.30 ± 2.63 g/L) (Figure 2B). Serum creatinine dosages confirmed severe kidney failure in these mice (135.58 ± 19.23 vs 7,89 ± 0.47 and 9,20 ± 1.33µM/L in 8 month-old WT and DH controls, respectively). No increase in serum creatinine was observed in healthy 8 month-old κF-DH mice (Figure 2C). At humane endpoints, macroscopic observations revealed pale and atrophic kidneys (Supplemental Figure 1) and some mice presented with pericardial and lung effusion (data not shown). Altogether, these results confirmed that the short life span of κF-DH mice is due to end-stage renal failure.

**Figure 2:**
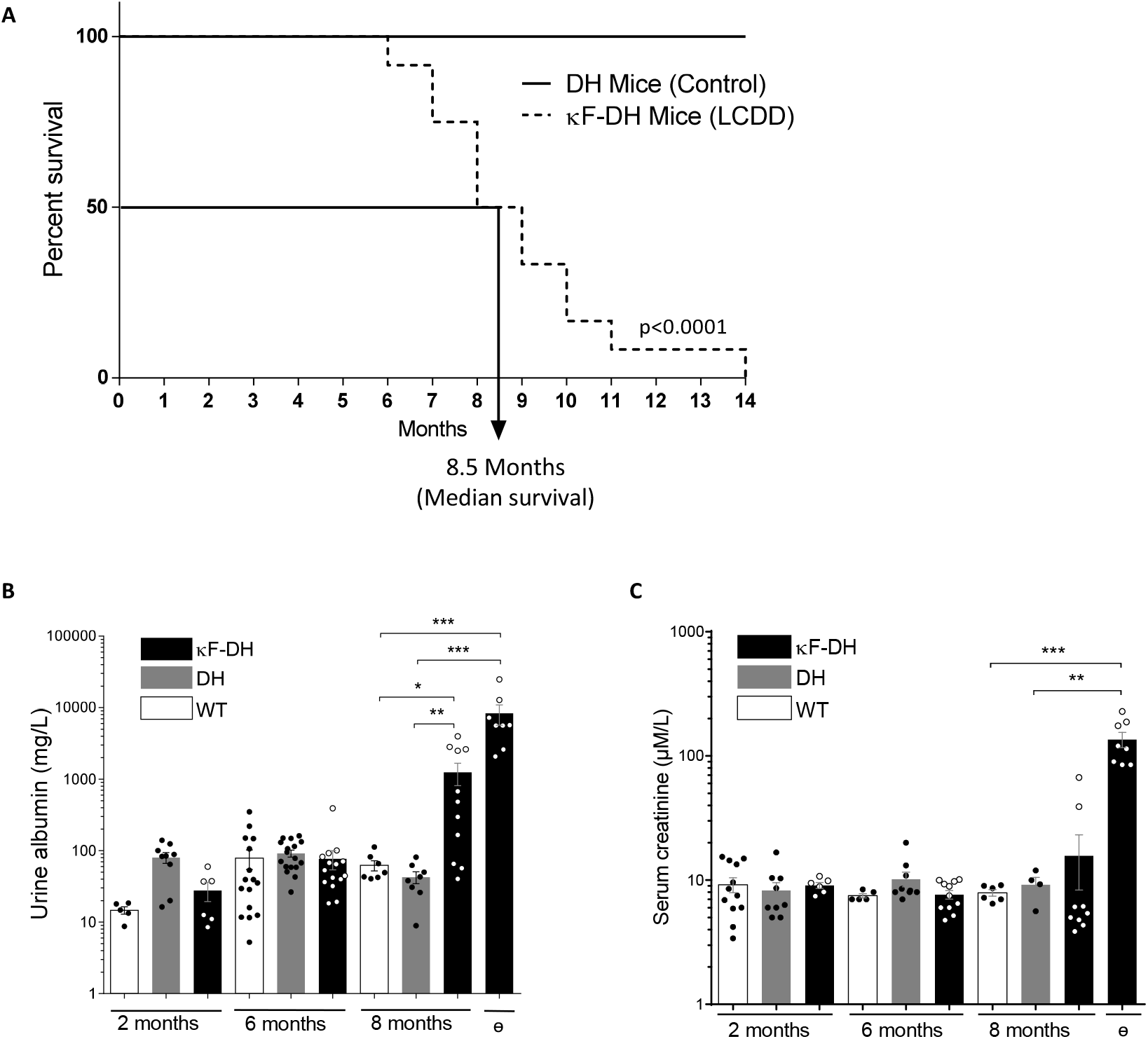
Survival analysis and kidney function in κF-DH mice. (A) Kaplan-Meier overall survival analysis of κF-DH mice compared to DH-mice. Note the short 8.5 months median survival for κF-DH mice compared to DH-mice (all alive at 14 months). (B) Urine albumin and (C) serum creatinine levels in 2 to 8-month-old κF-DH, DH and WT mice, and in κF-DH mice at humane endpoint (ɵ). Urine albumin in κF-DH mice started to increase at 8 months while a strong increase of serum creatinine appeared only in unhealthy mice. Survival data were analyzed using log-rank (Mantel-Cox) test and comparisons between 2 groups were calculated using the non-parametric Mann-Whitney test. Means are ± SEM and only significant *p* values are indicated. \**p* < 0.05; \*\**p* < 0.01; \*\*\**p* < 0.001; \*\*\*\**p* < 0.0001.

### Pathogenic human LCs recapitulate renal lesions of LCDD

Pathologic studies of kidney sections were carried out on 2- and 6-month-old κF-DH mice and in mice sacrificed at humane endpoints. As early as 2 months, immunofluorescence analysis revealed linear LC deposits along glomerular BMs (Figure 3A). BMs from medullary distal tubules were also readily positive for LC staining (Supplemental Figure 2). At 6 months of age, an intense LC staining was observed along all BMs in the renal medulla and cortex, and into mesangial space of the glomeruli (Figure 3A and Supplemental Figure 2). Despite abundant deposits and massive LC reabsorption in proximal tubules, most kidneys showed normal tubular and glomerular structures (Figure 3A and Supplemental Figure 2). Mice sacrificed at humane endpoints displayed absence of detectable LC reabsorption in proximal tubules, but massive staining of tubular BMs, of the mesangium and along Bowman’s capsule, with diffuse nodular appearance of glomerular deposits (Figure 3A and Supplemental Figure 2). At low magnification, corticomedullary dedifferentiation suggestive of severe renal failure was observed (Supplemental Figure 1). None of these staining patterns was observed in control mice. Periodic-acid Schiff staining confirmed diffuse thickening of tubular BMs, mesangial extracellular matrix expansion with mesangial cell proliferation and mesangiolysis, global nodular glomerulosclerosis with glomeruli enlargement (Figure 3B). Focal areas of interstitial fibrosis with inflammatory infiltrates were observed around sclerotic glomeruli and dilated tubules (Figure 3B). These infiltrates were mainly composed of granulocytes/macrophages (CD11b^+^) with few dendritic cells (CD11c^+^) and T cells (CD3^+^) (Supplemental Figure 3). Electron microscopy revealed characteristic linear “powdery punctuate” deposits on the inner aspect of glomerular BMs and the outer aspect of tubular BM, diffuse thickening of tubular and glomerular BMs, and expanded mesangial areas with massive accumulation of electron-dense material (Figure 3C). Finally, immunoelectron microscopy using gold-labelled anti-mouse kappa LC showed intense staining confirming the composition of the deposits (Figure 3D). Lesions of thrombotic microangiopathy with widening of the subendothelial zone by electron lucent material were seen in some glomeruli from 6 month-old mice and in most glomeruli from mice sacrificed at humane endpoints (Figure 3, C and D). Similarly to the patient from whom the LC gene was extracted (24), κF-DH mice also yielded LC deposits in other organs including lung, liver and heart (Supplemental Figure 4). Collectively, kidney lesions observed in κF-DH mice fully recapitulate the human pathological features of MIDD and are consistent with premature death due to end stage kidney failure.

**Figure 3:**
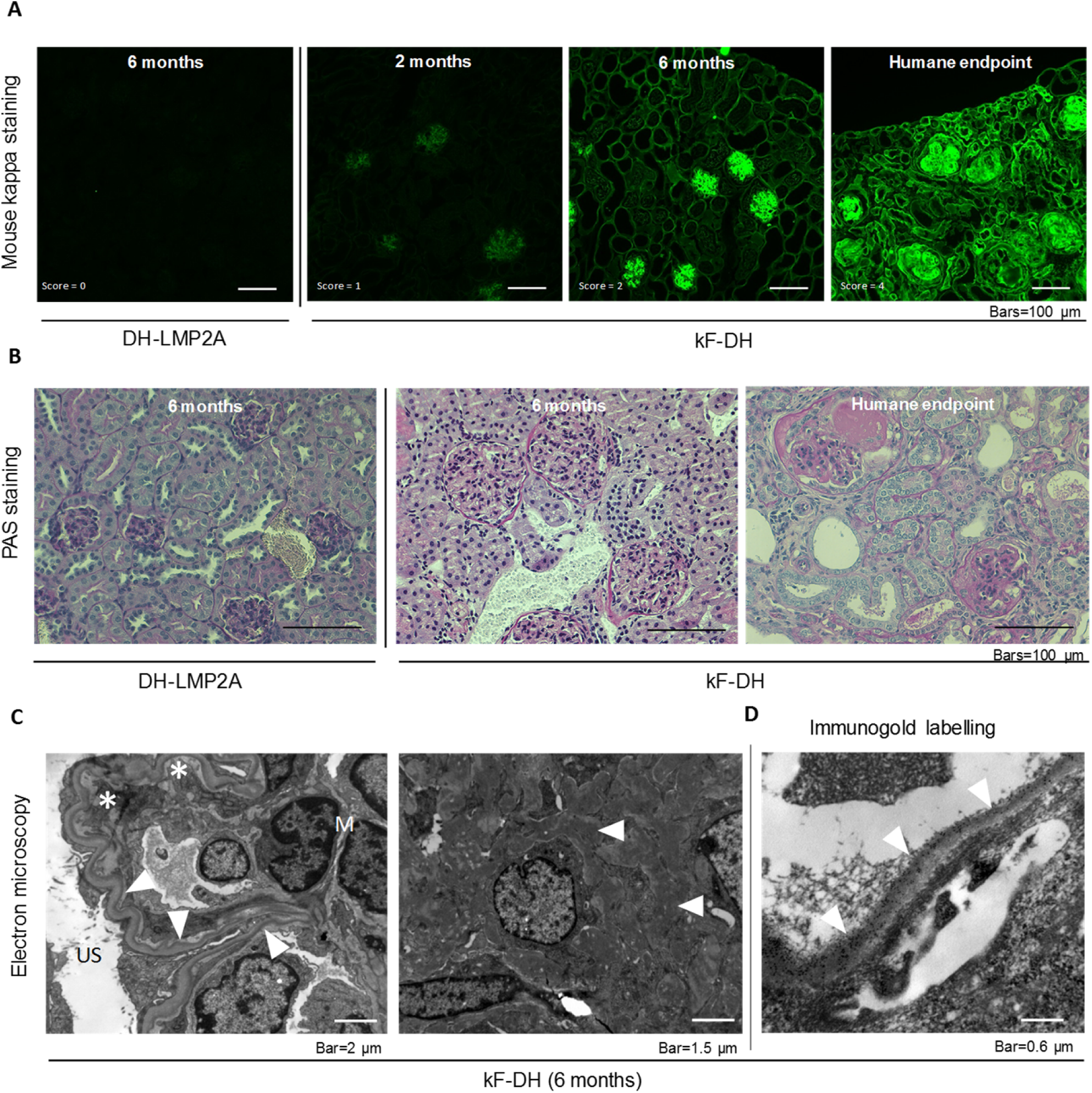
Kidney sections analysis of κF-DH mice. (A) Immunofluorescence microscopy on frozen kidney sections of κF-DH mice using an anti-mouse kappa antibody at 2 and 6 months and at humane endpoint, compared to DH control mice. Kappa light chain deposits in κF-DH are detectable as soon as the age of 2 months and become intense with age along tubular and glomerular basement membrane and in the mesangium. Lesion scores are indicated (see methods). Original magnification ×200. (B) Periodic acid Schiff (PAS) staining performed on paraffin embedded kidney sections showing tubular basement membranes thickening and nodular glomerulosclerosis with mesangial expansion of the extracellular matrix and mesangial cell proliferation. Note the massive enlargement of glomeruli in κF-DH mice. Original magnification ×200. (C) Electron microscopy analysis showing in the left panel linear electron-dense deposits in the inner aspect of the glomerular basement membrane (arrowhead). Subendothelial space is extremely widened consistent with thrombotic microangiopathy-like lesions (asterisk). Original magnification ×8,000. The right panel shows granular electron-dense deposits in the mesangium (arrow-head). Original magnification ×10,000. (D) Immunoelectron microscopy (original magnification ×40,000). Presence of anti-kappa gold-conjugated particles along the inner aspect of glomerular basement membrane (arrow-head). All data are representative of at least 3 kidneys in each group.

### Early induction of cell cycle and ECM remodelling are involved in LC-induced glomerulosclerosis

Having demonstrated that most 6-month-old κF-DH mice presented with early renal lesions of LCDD, including glomerular enlargement, mesangial cell proliferation and ECM expansion (Figure 3), we sought to compare their transcriptome to glomeruli from control mice. RNA-seq performed on purified glomeruli from 4 mice of each strain (and 5 WT) revealed a total of 360 and 475 genes expressed differentially in κF-DH mice compared to WT and DH controls, respectively (abs(log2FC) ≥ 0.7, FDR ≤ 0.05) (Figure 4A and Supplemental Figure 5A). Among these genes, 192 were deregulated compared to both controls, with 155 upregulated and 37 downregulated genes (Supplemental Figure 5A and B). We performed a C2 canonical pathway analysis from the Molecular Signatures Database (MSignDB v6.2) (25, 26) which revealed that the cell cycle (mitotic) was the main modified pathway (Figure 4B). Among the 20 more activated pathways (FDR *q* value from 5.93*10^−22^ to 1.78*10^−5^), 9 were directly related to the cell cycle (Figure 4B and Supplemental Table 1). Interestingly, all deregulated genes involved in the cell cycle were upregulated (Figure 4A, Supplemental Figure 5A and B and Supplemental Table 1). Deregulated genes in pathways not directly related to the cell cycle, like AURORA B, PLK1, FOXM1 or E2F, in fact frequently overlap with those of the cell cycle (Supplemental Table 1). Finally, beside the cell cycle, and as expected, we found 24 deregulated genes involved in the matrisome pathway (18 upregulated, 6 downregulated), including the previously described upregulation of Tenascin C (Tnc) and CTGF, known to be involved in glomerulosclerosis and interstitial fibrosis (Figure 4, A and B, supplemental Figure 5B and supplemental Table 1) (16, 27, 14). However, we did not find any activation of pathways related to TGFβ despite its established role in glomerulosclerosis. Immunostaining for Ki67 (Figure 4D) and Tenascin C (Supplemental Figure 6) in 6-month-old mice confirmed extensive cell cycle and ECM remodelling, respectively, in the glomeruli of κF-DH mice.

**Figure 4:**
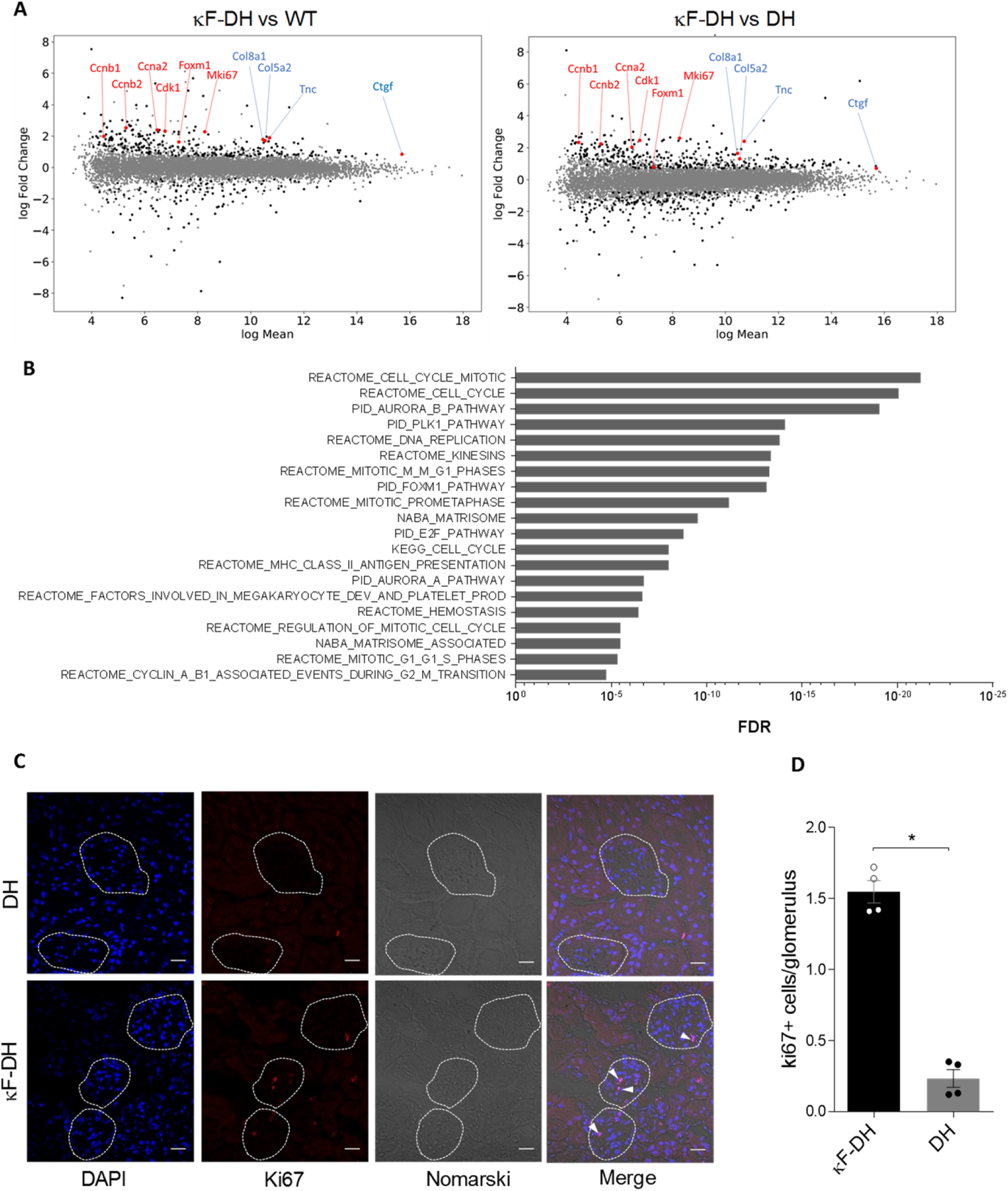
Glomerular transcriptomic analysis of κF-DH. (A) MA plots of normalized transcript values from κF-DH vs WT (left) and κF-DH vs DH (right) glomeruli. Examples of overexpressed genes in the cell cycle (red) and matrisome (blue) pathways are represented. (B) Top 20 enriched pathways based on the C2 canonical pathways from the Molecular signature database (MSignDB v6.2). All differentially expressed genes in κF-DH compared to both WT and DH glomeruli (see supplemental Figure 5A) are included in the analysis (155 upregulated and 37 downregulated genes). FDR *q*-value of the overlaps are represented on the bar graphs. (C) Intracellular Ki67 staining on frozen kidney sections of κF-DH compared to DH mice, the analysis was performed on 5 months-old mice. Glomeruli are surrounded and positive cells appear in red (arrowhead). (D) Mean of Ki67 positive cells in glomeruli of a complete kidney section of κF-DH compared to DH mice (n=4 of each strain). Note the increase of Ki67 positive cells in κF-DH glomeruli compared to DH mice. Means are ± SEM and comparisons between 2 groups were calculated using the non-parametric Mann-Whitney test. \**p* < 0.05

### LCDD LC induces ER stress and sensitizes plasma cell to proteasome inhibitors

Several studies tend to demonstrate that the production of a pathogenic monoclonal Ig or Ig fragment could influence plasma cell fitness and sensitivity to treatments through a general cellular stress involving ER stress, decreased autophagy and oxidative stress (17, 28). To determine if such cellular stress occurred in plasma cells producing LCDD LC and to better delineate the molecular pathways involved, we performed a RNA-sequencing (RNA-seq) on magnetically sorted spleen plasma cells (>80%) of κF-DH mice compared to WT mice producing polyclonal complete Ig, and to DH mice producing polyclonal FLCs (n = 3 mice of each strain). Our data revealed a total of 1946 and 809 expressed genes deregulated in LCDD LC-producing PCs compared to WT and DH controls respectively (abs(log2FC) ≥ 0.7, FDR ≤ 0.05). Among these deregulated genes, 722 and 376 were upregulated and 1224 and 433 were downregulated, respectively (Figure 5A). Because some deregulated genes in each comparison could result from differences between the WT and DH control PCs (22), only genes similarly deregulated in both comparisons (κF-DH *vs* WT and κF-DH *vs* DH) were considered for further analysis (Supplemental Figure 7A). This restricted the analysis to 180 and 213 genes, respectively upregulated and downregulated in κF-DH *vs* both controls (Supplemental Figure 7A and B). We performed a C2 canonical pathway analysis from the Molecular Signatures Database (MSignDB v6.2) and found that among the upregulated genes, the only significant pathways were related to the unfolded protein response (UPR) (Figure 5B and Supplemental Table 2). Genes of the UPR pathway upregulated in κF-DH PCs as compared to both controls are indicated on the MA-plots (Figure 5A) showing that they were not only upregulated but, as expected, were also highly expressed in PCs (29). Analyses using a classification of the differentially expressed genes by biological process (“GO biological process” from the Molecular Signatures Database) showed that the three most significant pathways were related to topologically incorrect proteins and ER stress, further suggesting that the activation of the UPR was likely due to the pathogenic LCDD LC (Figure 5C and Supplemental Table 3). Among the twenty most significant overlaps, nine were related to response to incorrect protein folding, ER stress or general response to stress. Downregulated genes were associated with ECM modification, hematopoietic lineage and G protein-coupled protein signalling (Supplemental Table 4). Gene set enrichment analyses (GSEA) on total genes confirmed that the UPR was significantly enriched in κF-DH PCs compared to both WT and DH cells (Supplemental Figure 7C). We performed real-time PCR on selected genes (Herpud1, Hspa5, Ddit3, Xbp1s) and confirmed a similar tendency for upregulation of genes involved in UPR even if, consistent with RNA-Seq analysis, no increase in Xbp1s was observed (Figure 5D). Having demonstrated that LCDD LC-producing PCs have an exacerbated UPR, we next verified if such ER stress could sensitize PCs to proteasome inhibitors (PI), as we previously showed in a HCDD mouse model (17). We treated mice with suboptimal doses of bortezomib (Bz) for two consecutive days to evaluate the level of PC depletion. This protocol led to a partial (about 2 fold) reduction of the absolute number of PCs in WT and DH mice (45.16 % and 57.54 % of depletion, respectively) as previously demonstrated (17). In contrast, PC depletion in κF-DH mice was far more efficient than in control mice with a mean depletion of 84.46 % corresponding to 6.44 fold reduction in absolute PC number (Figure 5E).

**Figure 5:**
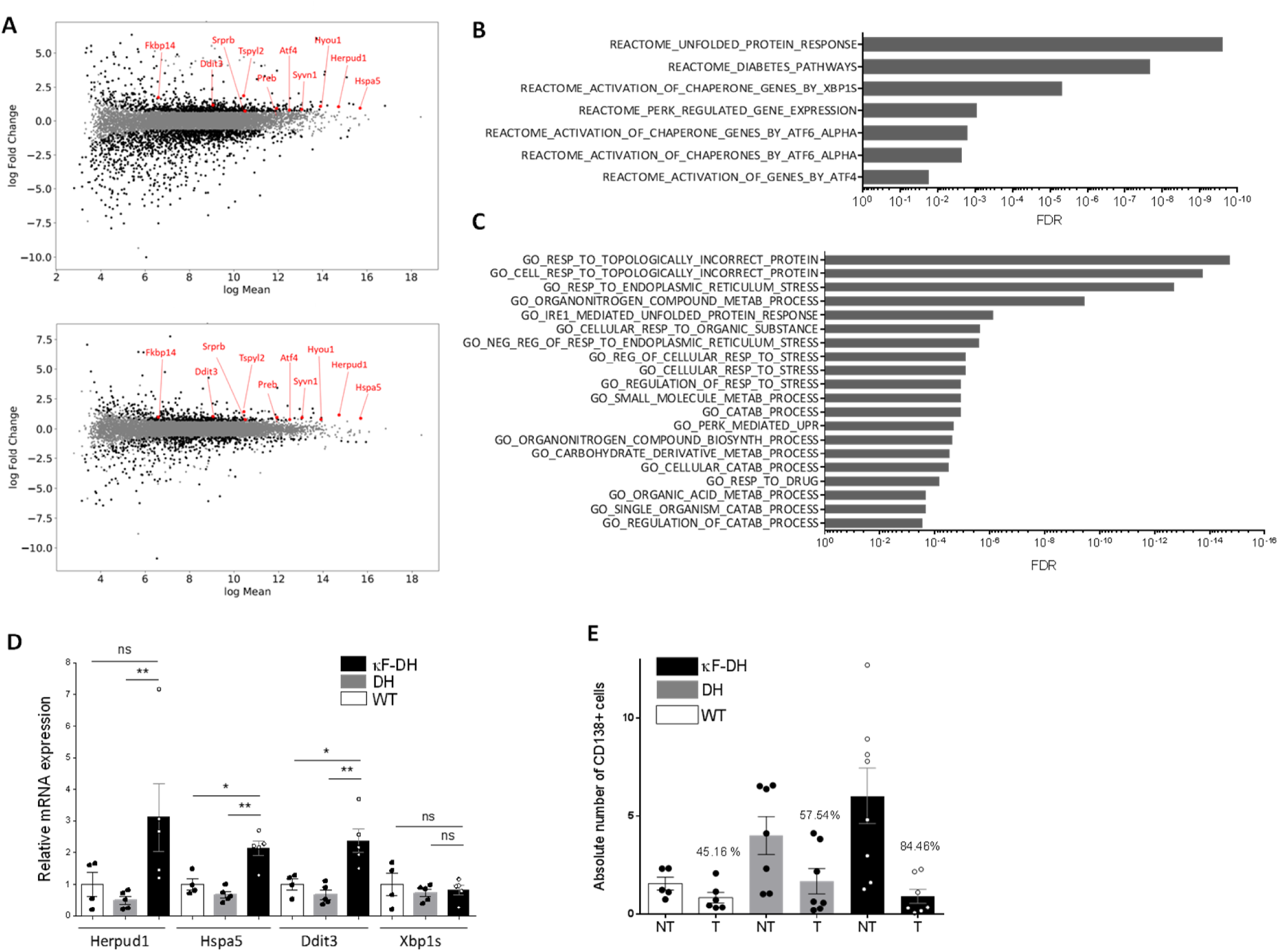
ER stress and sensitivity to proteasome inhibitors of κF-DH plasma cells. (A) MA plots of normalized transcript values from κF-DH vs WT (top) and κF-DH vs DH (bottom) plasma cells. Overexpressed genes, common in both comparisons, from the REACTOME UPR pathway are represented. (B) Significantly enriched pathways based on the C2 canonical pathways from the Molecular signature database (MSignDB v6.2). Analysis was done on the 180 up-regulated genes in κF-DH *vs* both controls (WT and DH PCs). FDR *q*-value of the overlaps are represented on the bar graphs. (C) GO biological process enrichment analysis. The enrichment was done on the 180 up-regulated genes in κF-DH *vs* both controls (WT and DH PCs). FDR *q*-value of the overlaps are represented on the bar graphs. (D) Quantitative transcriptional analysis of ER stress markers Herpud1, Hspa5, Ddit3 and Xbp1s in sorted CD138+ plasma cells of κF-DH, DH and WT mice. The increase of Herpud1, Hspa5, Ddit3 but not Xbp1s in κF-DH transcripts corroborates the transcriptomic analysis (n = 4 mice of each group) (E) Comparison of the absolute number of splenic CD138+ plasma cells between proteasome inhibitor treated (T) mice and non-treated (NT) mice. Percentage of depletion for each group is indicated over the bars (N = 5-8 mice of each group in 3 independent experiments). Means are ± SEM and comparisons between 2 groups were calculated using the non-parametric Mann-Whitney test. ns, non-significant; * *p* < 0.05; ** *p* < 0.01

### Removal of circulating LC leads to a rapid decrease of renal LC deposits and protects mice from kidney failure

The efficient response to PI prompted us to evaluate the effect of LC removal on renal lesions evolution. Weekly injections of Bz + cyclophosphamide led to an almost complete (> 10-fold reduction) and sustained depletion of circulating κ LC (Supplemental Figure 8). Treatment was administered for 2 months and all mice were subsequently sacrificed and processed for renal histological studies, except for mice reaching humane endpoints during the course of the experiment, which were immediately sacrificed. As shown in Figure 6A, treatment completely protected mice from early death. Kidneys from treated and non-treated mice were analysed for LC deposits and scores were established to compare the amount of lesions. As expected, treated mice presented significantly less abundant renal LC deposits as compared to non-treated mice (Figure 6 B and C). Interestingly, kidney lesions after treatment were also significantly lower as compared to non-treated 6-month-old mice (used in Figure 2 and 3), corresponding to the age of mice at start of the treatment (Figure 6 B and C). PAS staining showed glomeruli of smaller size as compared to 6-month-old mice and EM analysis revealed less marked glomerulosclerosis and the absence of thrombotic microangiopathy-like lesions, despite persistence of diffuse thickening of glomerular BMs (Figure 6 B and C). Biochemical studies confirmed that treatment efficiently halted the progression of kidney dysfunction. While serum creatinine was significantly increased in non-treated mice, it was maintained at normal levels in treated mice, which, compared to control 6-month-old mice, showed slight but non-significant increase in albuminuria (Figure 6E).

**Figure 6:**
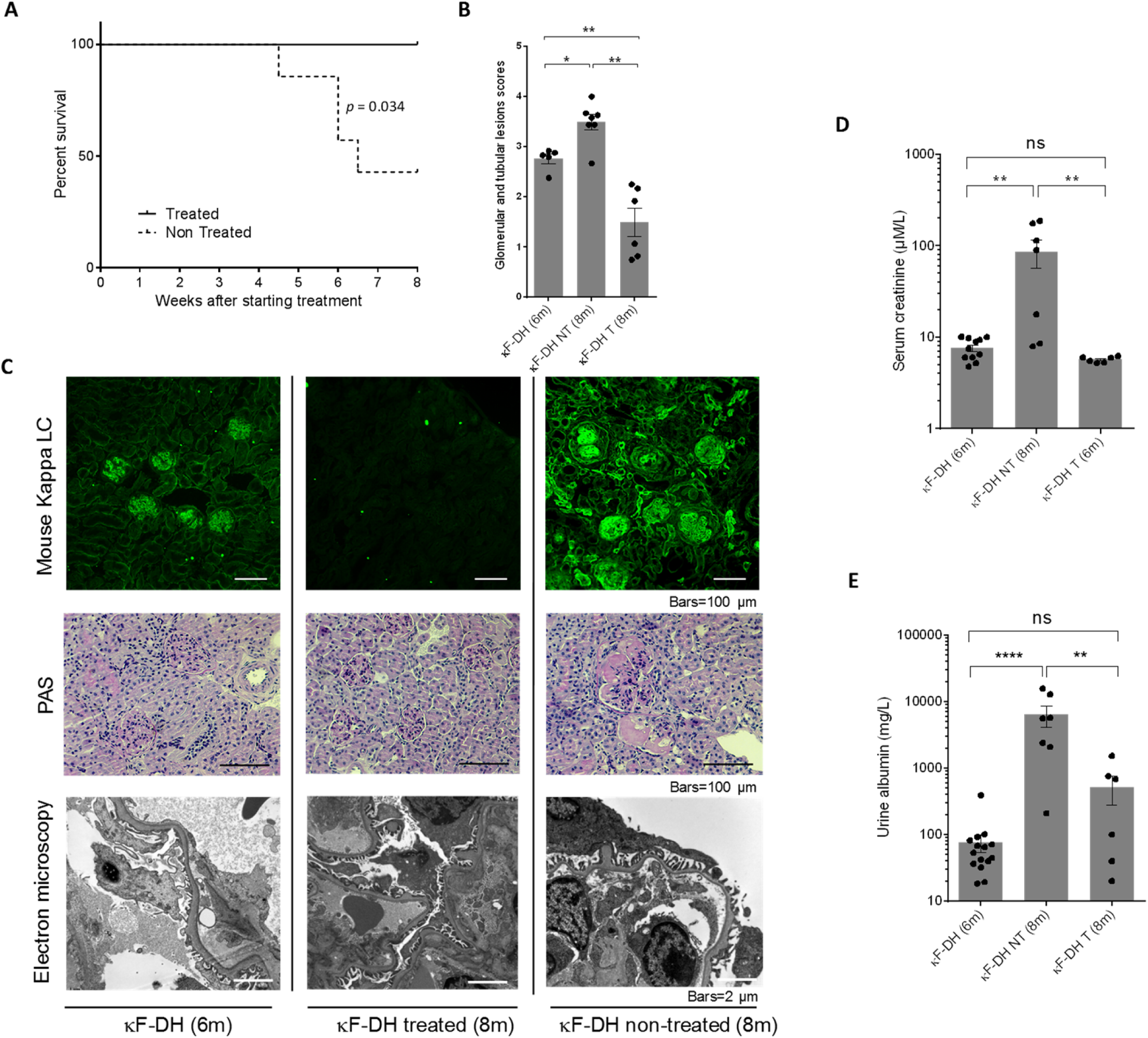
Effect of bortezomib/cyclophosphamide-based therapy on survival, renal lesions and function of κF-DH mice. (A) Kaplan-Meier overall survival analysis of non-treated (n=7) versus treated (n=6) κF-DH mice during the 8 weeks of treatment (from the age of 6 to 8 months). Note that none of the treated mice died during the curse of the treatment while half of the non-treated control mice died. Results are from 2 independent experiments (B) Comparison of kappa light chain deposits in kidney of κF-DH mice at the age of the beginning of the treatment (6-month-old κF-DH, n = 5) and the kidneys of treated (8-month-old κF-DH T, n = 6) and non-treated (8-month-old κF-DH NT, n = 7) mice at the end of the treatment. Evaluation of total deposits, glomerular and tubular, were performed with an anti-mouse kappa light chain staining by 4 different experimenters in n = 2 blinded experiments. (C) Lesions analysis on kidney sections by immunofluorescence microscopy (top), light microscopy (middle) and electron microscopy (bottom) between κF-DH mice at the beginning of the treatment and mice treated/non-treated at the end of the treatment. Note that deposits revealed by immunofluorescence in non-treated mice or mice at the beginning of the treatment totally disappeared in treated mice. LCDD suggestive lesions are also absent in treated mice (no glomerulosclerosis or thrombotic microangiopathy-like lesions). Original magnification ×200 (D and E) Serum creatinine and urine albumin levels in treated (κF-DH T) and non-treated (κF-DH NT) mice compared to six-month old mice (κF-DH). Note the absence of kidney dysfunction in treated mice. Survival data were analyzed using log-rank (Mantel-Cox) test and comparisons between 2 groups were calculated using the non-parametric Mann-Whitney test. Means are ± SEM and only significant *p* values are indicated. \**p* < 0.05; \*\**p* < 0.01; \*\*\**p* < 0.001; \*\*\*\**p* < 0.0001.

## Discussion

We herein described the first transgenic mouse model fully reproducing specific renal lesions and kidney dysfunction observed in human LCDD. We previously characterized another model of MIDD using a similar transgenic strategy, i.e. targeted insertion of a human pathogenic Ig gene into the mouse kappa locus (17). In this model, the human Ig gene was obtained from a patient with biopsy-proven HCDD and was composed of a truncated γ1 HC lacking the CH1 domain. Production of the pathogenic Ig in this model induced kidney lesions closely resembling those observed in human MIDD, including diffuse tubular and glomerular BM thickening and mesangial deposits. However, mice did not develop glomerulosclerosis, a hallmark of HCDD (5), nor exhibited glomerular dysfunction (normal creatinine and albuminuria). We suspected the genetic background to be involved, since C57BL6 and 129SV mice were previously shown to be highly resistant in models of nephron reduction or diabetic nephropathy (30–32). However, we could not exclude that the low level of circulating HC (~30-40 µg/ml) precluded pathological and clinical progression of the disease. In the present LCDD mouse model, we used a new strategy to obtain high levels of circulating human FLC (33, 18). We inserted the pathogenic V domain of a human LC into the kappa locus and crossed the mice with DH-LMP2A mice that display increased plasma cell development in the absence of Ig heavy chains (21, 22, 18). Using this approach, production of a chimeric FLC (composed of a human pathogenic Vκ4 domain and the mouse Cκ domain) occurred in virtually all plasma cells and kappa LC serum levels reached more than 1g/l in κF-DH mice, which is similar to that observed in LCDD patients (23). Similar productions were obtained in other unpublished models of MGRS (18), making this strategy a powerful tool to study the pathogenicity of monoclonal LCs *in vivo*.

Accordingly, κF-DH mice recapitulate progressive glomerular lesions closely resembling those observed in MIDD patients, with linear monoclonal kappa LC deposits along glomerular and tubular BMs and in the mesangium, with glomerular enlargement and mesangial hyper cellularity, resulting in albuminuria. The lesions eventually lead to nodular glomerulosclerosis and end-stage kidney failure within the first year. Then we first confirmed that the V domain bears alone the pathogenic properties of LCCD LCs, similarly to what we shown in a model of LC-induced Fanconi syndrome (19). If the pathological lesions slowly progress during the first months, the occurrence of kidney dysfunction with increased serum creatinine, appears to be more sudden and unpredictable, requiring careful follow-up of mice after the sixth month. All these features make the κF-DH mice an accurate model to study the pathophysiology of MIDD, but also other of other kidney diseases featuring nodular glomerulosclerosis, particularly diabetic nephropathy (DN). Indeed, κF-DH mice recapitulate all the criteria used to validate the glomerular lesions of DN as established by the Diabetic Complications Consortium (34) and could be advantageously used to decipher common pathways involved in mesangial cell proliferation and ECM expansion leading to glomerulosclerosis. Nodular glomerulosclerosis has been widely studied, especially in the context of DN (35, 34, 36) and LCDD (37, 12, 15), highlighting a role for TGFβ signalling and PDGF. However, due to the lack of relevant animal models reproducing the chronic progression of kidney lesions, it is still unclear whether TGFβ and PDGF are early mediators of glomerulosclerosis or if they are involved latter, enhancing and accelerating terminal fibrosis of the glomeruli. When we performed RNA sequencing on glomeruli from 6-month-old LCDD mice which display abundant LC deposits but no glomerulosclerosis, we did not find any activation of the TGFβ or PDGF pathways despite already present ECM deregulation, as indicated by the overexpression of Tenascin C (16, 14, 38). However, we observed a slight but significant increase of CTGF, known for its pro-fibrotic properties in damaged tissues including the kidney (27, 39). The most prominent early effect of LC deposition in our model was the induction of cell cycle, as observed by RNA-seq and further confirmed by Ki67 staining. This result is consistent with the presence of hypercellularity and enlargement of glomeruli that precede glomerulosclerosis and kidney dysfunction. Similar studies in mice with more advanced disease stages deserve to be carried out to decipher the sequential molecular events leading to glomerulosclerosis, but our results suggest that TGFβ overproduction and its consequences are secondary events following LC-induced proliferation of mesangial cells. Understanding the pathways linking LC deposition and cellular proliferation should likely give new insights in the pathophysiology of the disease and should pave new ways for early therapeutic interventions.

Recent studies have highlighted the efficiency of PI-based treatment in patients with MIDD, as already demonstrated in AL-amyloidosis (40–43, 23, 5). We confirmed this observation in our mouse model using weekly injections of bortezomib plus cyclophosphamide for up to two months. This regimen led to a rapid hematological response with almost complete and sustained depletion of circulating FLC that was accompanied by a significant improvement in kidney lesions with decrease of glomerular and tubular LC staining. Interestingly, kidney lesions were significantly decreased compared to those observed in animals at age corresponding to the beginning of treatment, highlighting a possible “cleaning” mechanism leading to the elimination of the LC deposits. This improvement in kidney lesions was also characterized by normalization of the glomerular size and the absence of thrombotic microangiopathy-like lesions occasionally observed in six-month-old mice. Although complete recovery was not achieved after 2 months as shown by electron microscopy showing persistent thickened basement membranes and deposits in the mesangium, the treatment was sufficient to maintain normal serum creatinine levels with a slight increase in albuminuria. This result is in accordance with studies of human diseases, including diabetic nephropathy (44, 45) which showed that glomerulosclerosis at early stages can be reversed with efficient treatment, although this process in humans can take several years. In mouse models, however, the reversibility was demonstrated after short term treatment in diabetic nephropathy (46, 47) or hypertension-related glomerulosclerosis (48). Accordingly, we demonstrated the almost complete reversibility of kidney lesions in LCDD-induced glomerulosclerosis following LC depletion. Further studies are needed to confirm if long-term LC removal could lead to complete renal recovery, to decipher the involved mechanisms and to test therapeutic strategies that could accelerate this process. In this latter view, it seems that therapy based on mesenchymal stem cells could be a valuable approach, as demonstrated in rodent models of glomerulopathies (49, 50) or more recently in *ex vivo* models of monoclonal LC-induced glomerular damages (51). Our model also enabled us to analyse the intrinsic toxicity of pathogenic monoclonal LC for plasma cells. Similarly to what was shown in patients with MIDD (52, 23, 5, 10) and in the HCDD model (17), plasma cells producing LCDD LC are highly sensitive to PI. We assume that mouse plasma cells are quite different from dysplasic human plasma cells, in that genomic alterations other than the sole LC production could explain such sensitivity. However, on the contrary, differences in PI sensitivity in our model uniquely depends on the presence of the human LC since plasma cells are otherwise strictly identical. Using RNA-seq and real time PCR, we showed that these plasma cells display a high ER stress and more generally, an overexpression of genes involved in the response to misfolded or unfolded proteins. Similar results have been obtained with plasma cells forced to express heavy chain only (53), truncated HC (17) or truncated LC (54). Collectively, we hypothesize that the high sensitivity to proteasome inhibitors is likely related to the impaired capacity of plasma cells to cope with the production of an abnormal, aggregation-prone, monoclonal LC, as observed in AL amyloidosis (28). Using our transgenic strategy, we now have developed several mouse models expressing human LC from other MGRS, including AL amyloidosis and Fanconi syndrome (18), which will serve to determine if all pathogenic LC are toxic for plasma cells and to decipher the common pathways leading to ER stress and PI sensitivity.

We describe here the first mouse model fully recapitulating the features of MIDD, including glomeruloslerosis and end stage kidney disease. Compared to other models of chronic kidney diseases, the development of kidney dysfunction relies on the natural causing factor (*i.e.* monoclonal LC) without any chemical or physical induction or genetic mutations artificially triggering the disease (55). Consequently, such a model will likely be useful to understand the molecular mechanisms leading to glomerulosclerosis in MIDD but also other chronic kidney diseases such as diabetic nephropathy. It also confirms the toxicity of abnormal LC for plasma cells and will likely serve to design and test new therapeutic strategies for MIDD and other related diseases.

## Methods

### Mice

Gene targeting into the murine Igκ locus was performed as previously described (19). Briefly, the gene coding the variable region (VJ) of a human monoclonal κ light chain extracted from a patient with LCDD was introduced in place of the mouse Jκ segments. The procedure is detailed in the supplemental Methods. DH-LMP2A mice (21) were kindly provided by S. Casola (IFOM, Milan, Italy). All animals are under a mixed genetic background (BALB/C × 129SV × C57/BL6) and were maintained in pathogen-free conditions and analyzed at 2, 6 and 8 months except when otherwise stated. All experimental procedures have received the approval of our institutional review board for animal experimentation and of the French Ministry of Research (N° 7655-2016112211028184). Humane endpoints include lethargy, hunched posture, lack of response to manipulation, rough hair and visible weight loss. All mice starting to present one of the humane endpoint above were immediately sacrificed on a continuous-flow basis.

### *In vivo* treatment, surgery and biochemical parameters

Mice were treated with Bortezomib 0,75mg/kg (56) (Velcade^®^, Janssen Cilag) and cyclophosphamide (Endoxan^®^, Baxter) (2mg/kg) once a week during 2 months. Bortezomib was injected subcutaneously and cyclophosphamide was injected intraperitoneally. To test the sensitivity of plasma cells to Bortezomib, we daily injected a sub-optimal dose of Bortezomib for two days as previously described (17). All injections were performed under anesthesia. Biochemical parameters were measured on overnight urine collection and blood samples were obtained by retro-orbital puncture under anesthesia. Serum concentrations of creatinine were measured on a Konelab 30 analyser with a creatinine enzymatic test (ThermoFisher Scientific). Urine albumin concentrations were measured using an albumin mouse ELISA kit (Abcam), according to the manufacturer’s recommendations.

### Glomeruli extraction

Isolation of kidney glomeruli from mice was derived from Takemoto and al. (57). Briefly, mice were perfused in the heart with magnetic 4.5µm diameter Dynabeads. Kidneys were minced, digested by Dnase I (Roche) and collagenase IV (Sigma-Aldrich), filtered, and glomeruli were isolated using a magnetic rack. Total RNA from purified glomeruli were obtained using miRNeasy Mini kit (Quiagen).

### Flow cytometry

Intracellular staining was performed using the Intraprep™ kit (Beckman Coulter). Flow cytometry analysis were performed on a BD Pharmingen LSRFortessa^®^ cytometer. Data were analyzed with BD FACSDiva software (BD Biosciences). Antibodies used are listed in the Supplemental Table 5

### Western Blot

Serum proteins were separated by reducing SDS-PAGE (10%) and transferred onto polyvinylidene difluoride membranes (Millipore). Membranes were blocked in 5% milk Tris-buffered saline (TBS), then incubated with appropriate antibodies in 3% milk TBS (supplemental Table 2), washed three times with TBS 0.1 % tween and revealed by chemiluminescence (ECL, Pierce).

### Pathologic studies

Kidney samples were processed for light microscopic examination, immunofluorescence and electron microscopic studies, as previously described (19, 58). Briefly, immunofluorescence was performed on organs included in OCT and snap frozen in isopentane using a Snap Frost 2 (Excilone). Cryosections of 8µm were fixed with cold acetone, blocked with PBS 3% BSA and stained with appropriate antibodies (Supplemental Table 5). For ki-67 intracellular staining, fixed slides were permeabilized with PBS-tween 0.1 % and then stained in PBS-tween overnight. Total positive glomerular cells were counted manually on a complete kidney section. Slides were observed on a NiE microscope (Nikon). Immunoelectron microscopy was made on samples fixed with 4% glutaraldehyde in PBS and embedded in resin (TAAB Labs). Ultra-thin sections were processed for EM studies, incubated with anti-κ gold-conjugated and examined with a JEOL JEM-1010 electron microscope as previously described (59). Periodic acid schiff (PAS) stainings were prepared on paraffin-embedded kidney sections and examined by light microscopy on a NiE microscope (Nikon). Lesions scores were quantified on renal sections in blind experiments carried out by at least three independent experimenters. Lesions scores ranged from 0 to 4 (0 =absent; 1 = weak deposits on glomerular and tubular basement membranes; 2 = Bright deposits GBM and TBM but normal structures (absence of sclerosis); 3 = Bright deposits on GBM and TBM, start of sclerosis and abnormal structures in some tubules and glomeruli; 4 = Heavy deposits on GBM and TBM and glomerular sclerosis). Representative images with scores are depicted in Figure 3.

### In vitro stimulations

Spleen B cells were isolated and stimulated in vitro (1×10^6^ cell/mL) with 1µg/mL of LPS (InVivoGen) for four days, in DMEM supplemented with 10% FCS. 1.10^6^ cells were used for flow cytometry analysis.

### ELISA

Serum were analyzed for the presence of kappa light chains, as previously described (58). Plates were read at 405nm with a Multiskan FC® spectrophotometer (Thermo scientific).

### Transcriptional analysis

Total RNA was extracted using TRI Reagent (Ambion). After digestion of the remaining DNA with Dnase I (Invitrogen) on 1µg of total RNA, reverse transcription was performed using high capacity cDNA reverse transcription kit (Applied Biosystems), with random hexamers. Relative quantification was performed with Premix Ex Taq or SYBR Premix Ex Taq (Takara) on cDNA samples (10 ng per reaction). Quantification of the gene of interest was analyzed by ΔCt method with IgJ used as housekeeper gene. TaqMan^®^ probes for CHOP (Mm01135937_g1), BiP (Mm01333324_g1), IgJ (Mm00461780_m1) and a Sybr green^®^ assay for Xbp1-s (sXBP1for: 5’-gagtccgcagcaggtg-3’ and sXBP1rev 5’-gtgtcagagtccatggga-3’)(60) were used.

### RNA sequencing

Libraries were generated from 500 ng of total RNA using Truseq Stranded mRNA kit (Illumina). Libraries were then quantified with Qubit ds DNA HS assay kit (Invitrogen) and pooled. 4nM of this pool were loaded on a Nextseq 500 high output flowcell and sequenced on a NextSeq 500 platform (Illumina) with 2 × 75bp paired-end chemistry. Mapping was performed with STAR_2.4.0a versus mm10 following Encode RNA-seq options – outFilterIntronMotifs RemoveNoncanonicalUnannotated –alignMatesGapMax 1000000 – outReadsUnmapped Fastx –alignIntronMin 20 –alignIntronMax 1000000 – alignSJoverhangMin 8 –alignSJDBoverhangMin 1 –outFilterMultimapNmax 20 then a read counts table was generated with featureCounts (subread-1.5.0-p3-Linux-x86_64) with – primary -g gene_name -p -s 1 -M -C options. Quality control of RNAseq count data was assessed using in-house R scripts. Low expressed genes were filtered out, then normalization and statistical analysis were performed using Bioconductor package DESeq2. Further filtrations excluded genes from the Y chromosome, non-coding RNA, unknown transcripts, predicted genes (including Rik and Gm RNA) and pseudogenes. Candidate differentially expressed genes were selected using a combination of FDR < 0.05 and abs(log2FC) > 0.7. Venn diagrams were created using Venny 2.1 (http://bioinfogp.cnb.csic.es/tools/venny/index.html). Overlaps of differentially expressed genes with C2 Canonical Pathways and GO Biological Process gene sets from Molecular Signature Database v6.2 were obtained on the Gene Set Enrichment Analysis (GSEA) website (25, 26). Only the resulting top twenty gene sets from significant list (FDR < 0.05) were considered. Gene Set Enrichment Analysis of the REACTOME UPR gene set was done using the GSEA software (25). Gene Expression Omnibus (GEO) accession numbers for the RNAseq datasets reported in this paper are: GSE119049 for glomeruli and GSE119048 for plasma cells. All datasets can be found in the GEO superseries: GSE119050.

### Statistical analysis

The statistical tests used to evaluate differences between variables were carried out using Prism GraphPad software (GraphPad Software, Inc). Statistical significance were determined using the non-parametric Mann-Whitney test to assess two independent groups and the log-rank (Mantel-Cox) test for survival analyses. *p* values < 0.05 were considered significant.

## Author contributions

Contribution: S.B. designed, performed, and analyzed experiments and drafted the manuscript; A.B. designed, performed, and analyzed some experiments; M.V.A performed, and analyzed experiments and drafted the manuscript; V.J. drafted parts of the manuscript and provided general advice; A.R., N.Q., S.K., C.C., C.O., Z.O. and M.O.A performed and analyzed some experiments; F.B., B.H., A.P. and N.P. performed the biostatistics and bioinformatics on RNA-seq data; G.T. analyzed and provided advices on kidney pathology; A.J., L.D., M.C. provided general advice and reviewed the manuscript; F.B. analyzed data and critically reviewed the manuscript; C.S. designed and supervised research and wrote the manuscript.

## Acknowledgments

The authors thank the staff of the BISCEm technical platforms at the University of Limoges (Animal facility, cell cytometry, microscopy, transgenesis and bioinformatics facilities), the Department of Pathology of Poitiers, L. Magnol and K. Vuillier for access to the Konelab 30 analyser and A. Garot for helpful discussions. This work was supported by grants from Fondation Française pour la Recherche contre le Myélome et les Gammapathies monoclonales, Limousin committees of Ligue nationale contre le cancer, Fondation pour la Recherche Médicale, Agence régionale de la santé and Institut Universitaire de France. S.B. is supported by Centre Hospitalier Universitaire Dupuytren Limoges and Plan National Maladies Rares. M.V.A., A.B. and M.O.A were funded by fellowships from Région Limousin (now Région Nouvelle Aquitaine), French ministry of research and Agence de Valorisation de la recherche de l’université de Limoges.

