## Supplemental figures for "Glomerulosclerosis and kidney failure in a mouse model of monoclonal immunoglobulin light-chain deposition disease"

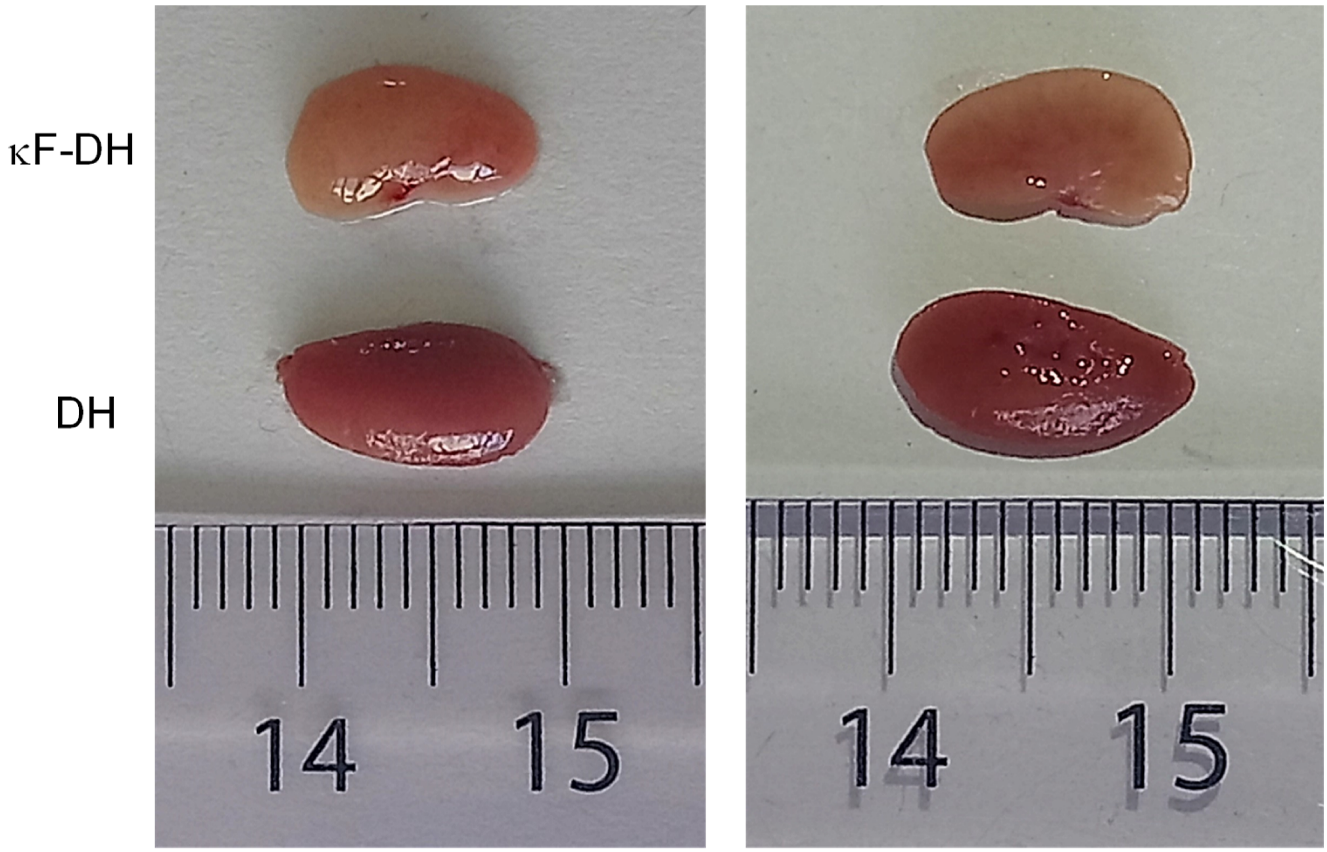


Supplemental Figure 1 : ***Ex-vivo* observations of kidneys from κF-DH vs DH-LMP2A mice.** Macroscopic observation revealed bloodless kidneys in κF-DH mouse at sacrifice (humane endpoint) compared to a DH mouse of the same age. Left panel: whole kidney, right panel: longitudinal section. Scale is in cm.

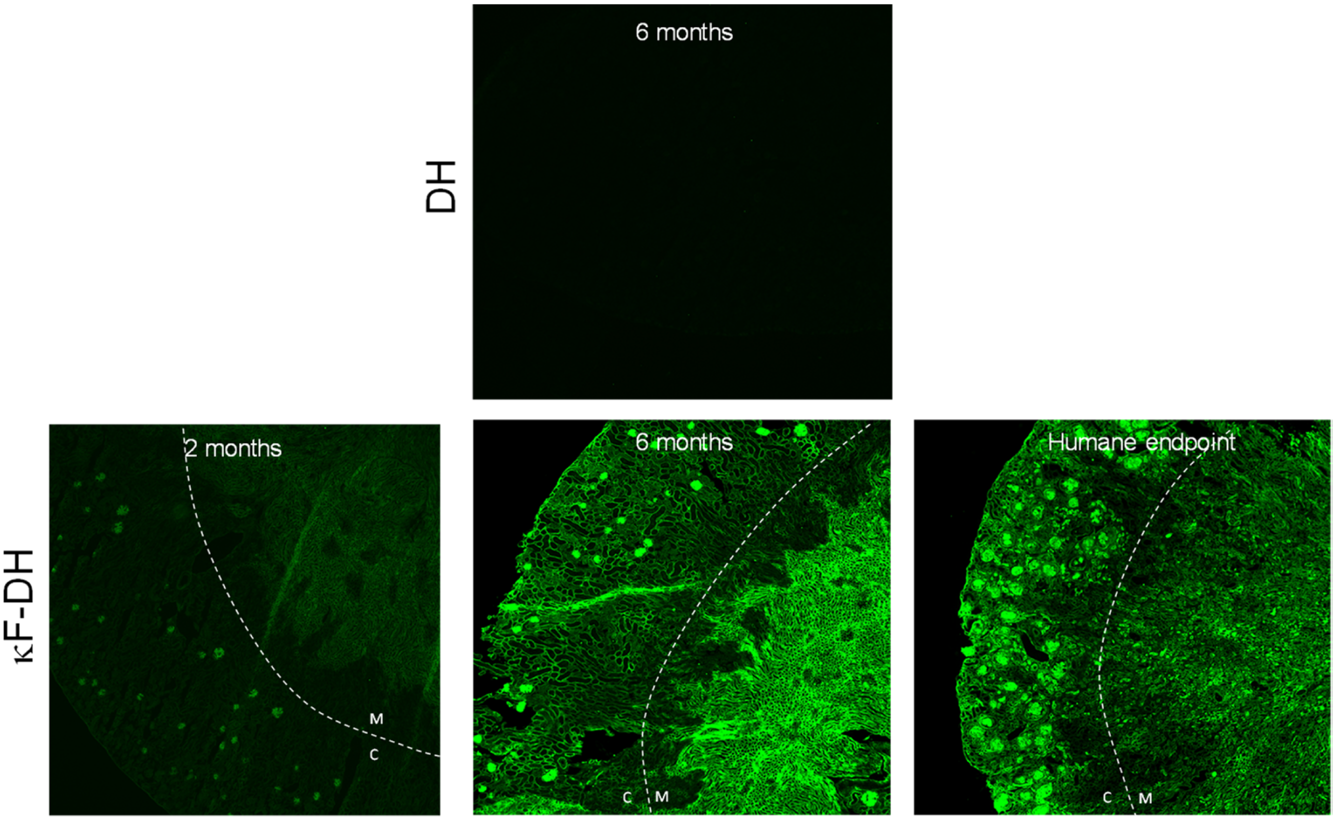


Supplemental Figure 2 : **Kidney sections analysis of κF-DH mice.** Immunofluorescence microscopy on frozen kidney sections stained with an anti-mouse kappa antibody of κF-DH mice at 2, 6 months and humane endpoint, compared to a DH control mouse. Kappa light chain deposits in κF-DH are detectable in the cortex (C) and medulla (M) of the kidney as soon as the age of 2 months. Original magnification x40.

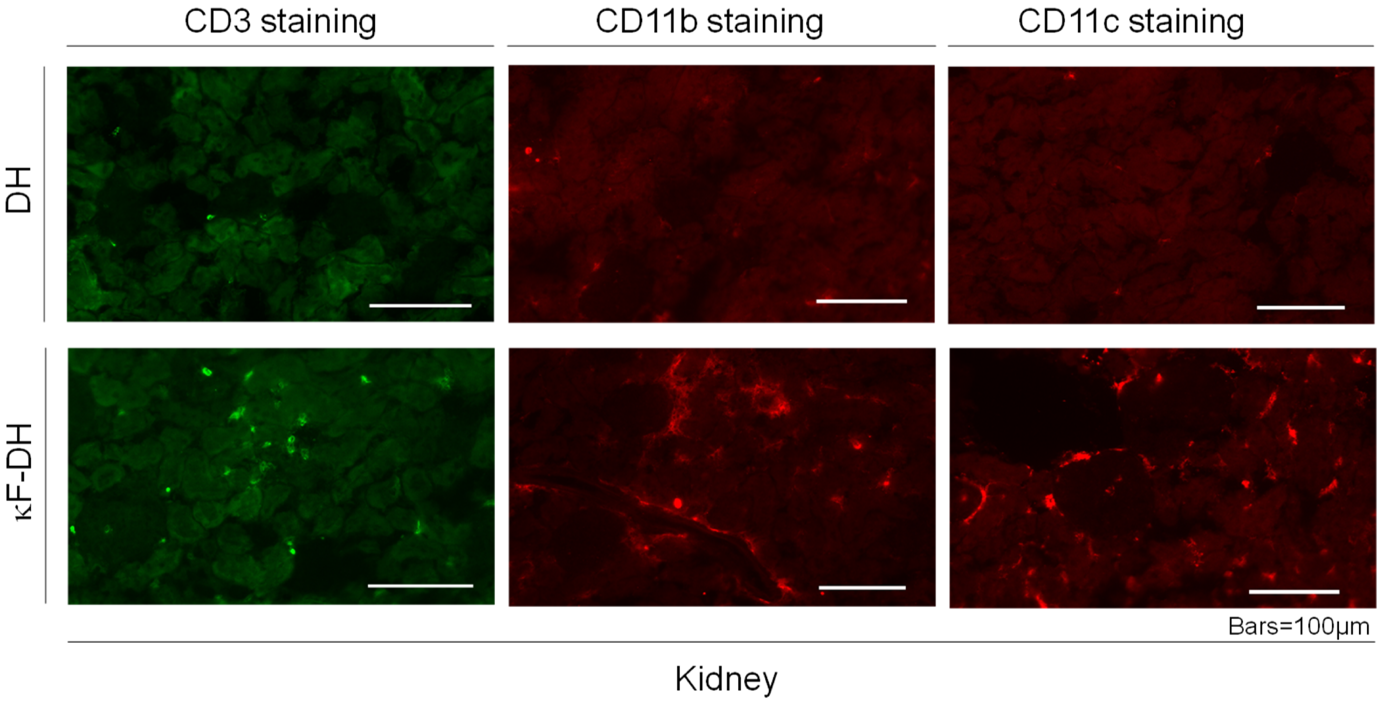


Supplemental Figure 3 : **Immune cells infiltrate the kidney of κF-DH.** Immunofluorescence microscopy on frozen kidney sections of κF-DH mice using an anti-mouse CD3 (T cells), CD11b (granulocytes/macrophages) or CD11c (dendritic cells) at 6 months compared to DH control mice. Original magnification x200.

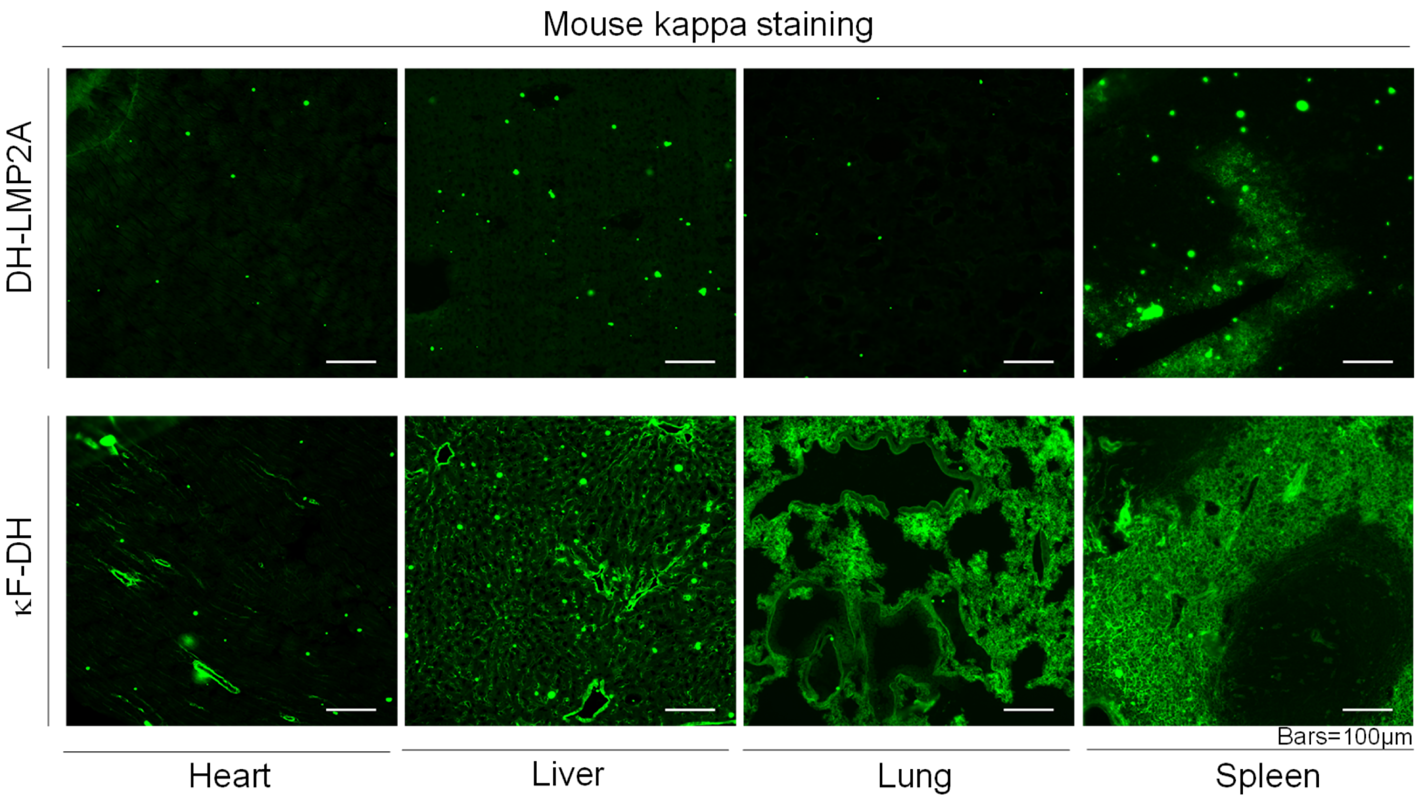


Supplemental Figure 4 : **Deposits detection in other organs of κF-DH mice.** Immunofluorescence microscopy on frozen heart, liver, lung and spleen sections of κF-DH mice using an anti-mouse kappa antibody at 6 months, compared to DH control mice. Kappa light chain deposits in κF-DH are detectable in all these organs. Original magnification x200.

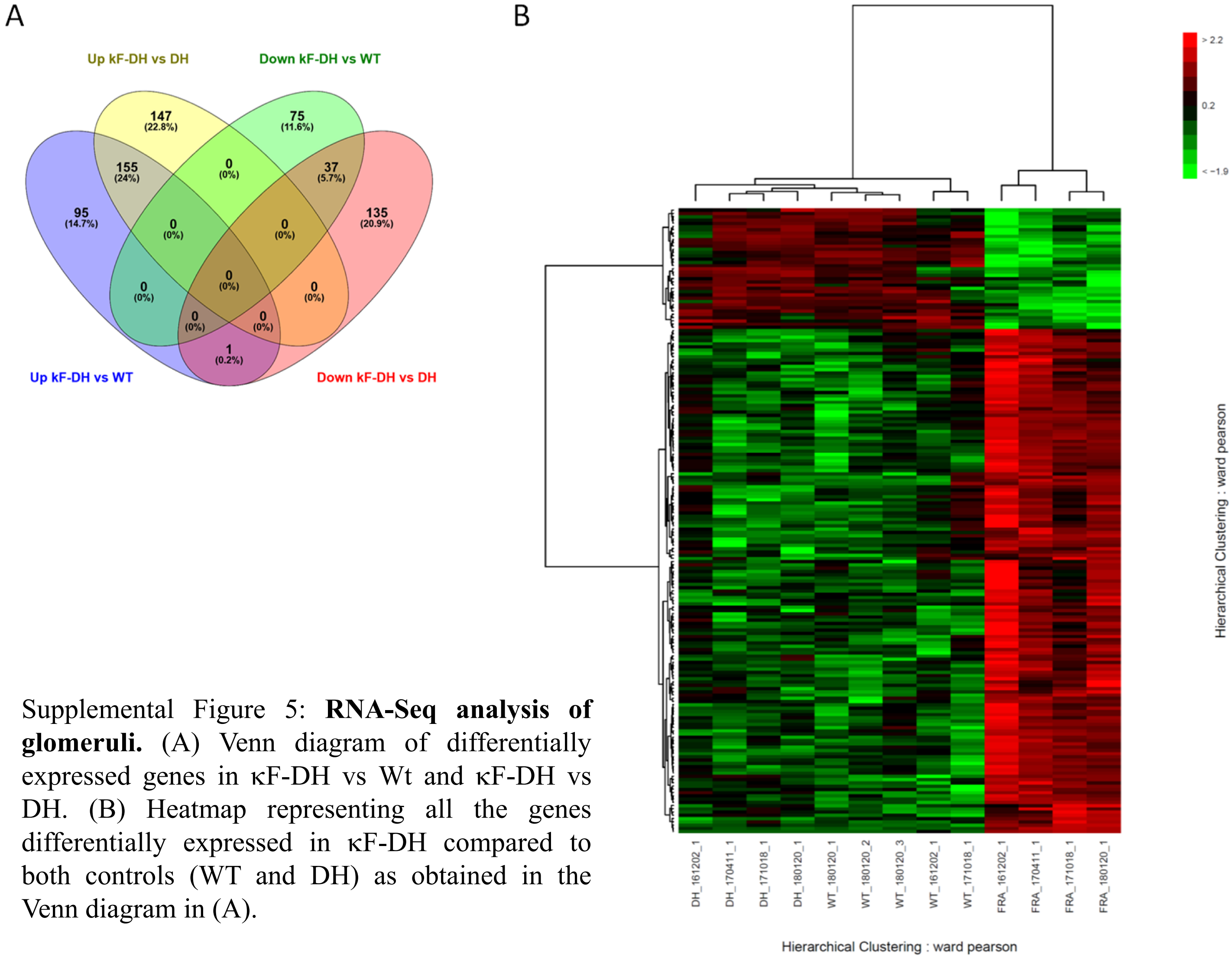


Supplemental Figure 5: **RNA-Seq analysis of glomeruli.** (A) Venn diagram of differentially expressed genes in κF-DH vs Wt and κF-DH vs DH. (B) Heatmap representing all the genes differentially expressed in κF-DH compared to both controls (WT and DH) as obtained in the Venn diagram in (A).

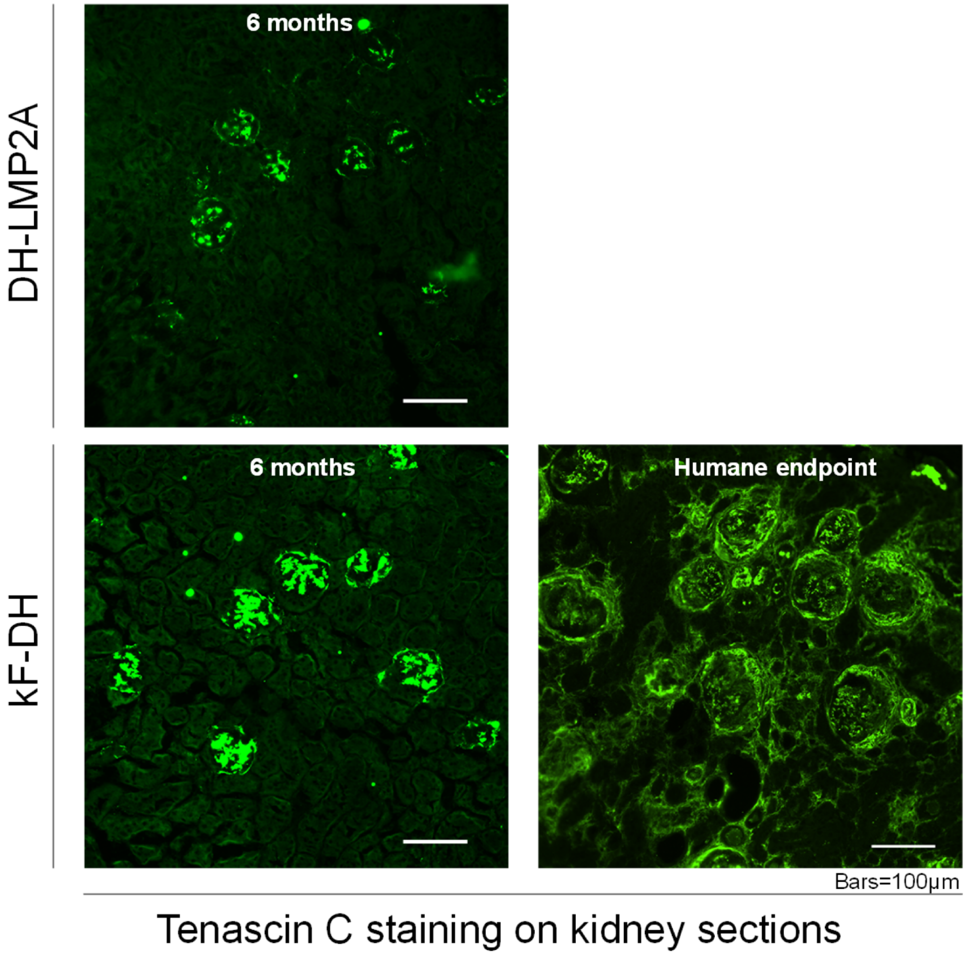


Supplemental Figure 6 : **Tenascin C analysis in kF-DH mice.** Immunofluorescence microscopy on complete frozen kidney sections of kF-DH mice using an anti-mouse tenascin C antibody at 6 months and mice at humane endpoint, compared to DH control mice. Original magnification x200.

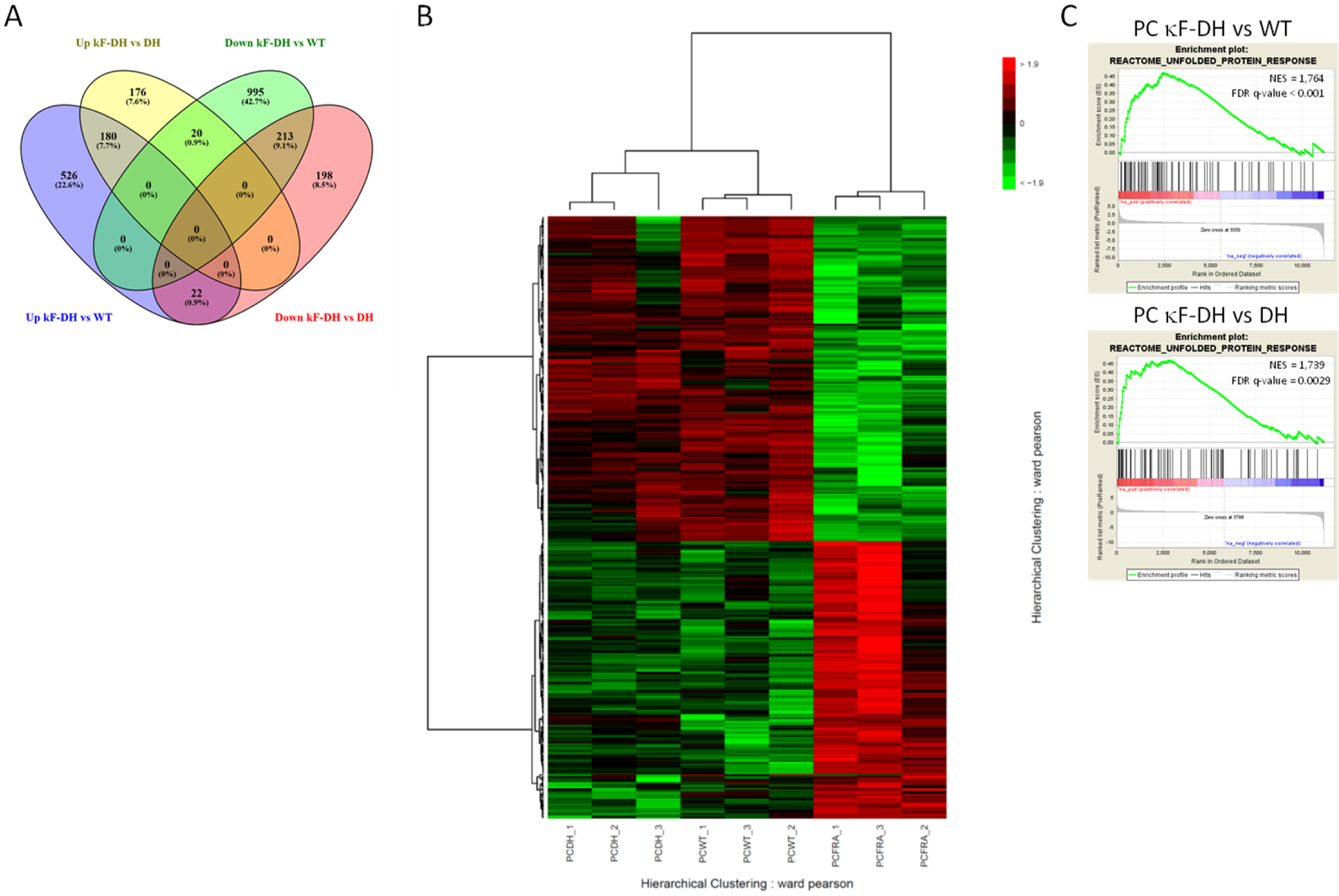


Supplemental Figure 7: **RNA-Seq analysis of plasma cells.** (A) Venn diagram of differentially expressed genes in κF-DH vs WT and κF-DH vs DH. (B) Heatmap representing all the genes differentially expressed in κF-DH compared to both controls (WT and DH) as obtained in the Venn diagram in (A). (C) Gene set enrichment analysis (GSEA) plots showing enrichment in the Reactome UPR pathway of κF-DH plasma cells compared to WT (Top) and DH (Bottom).

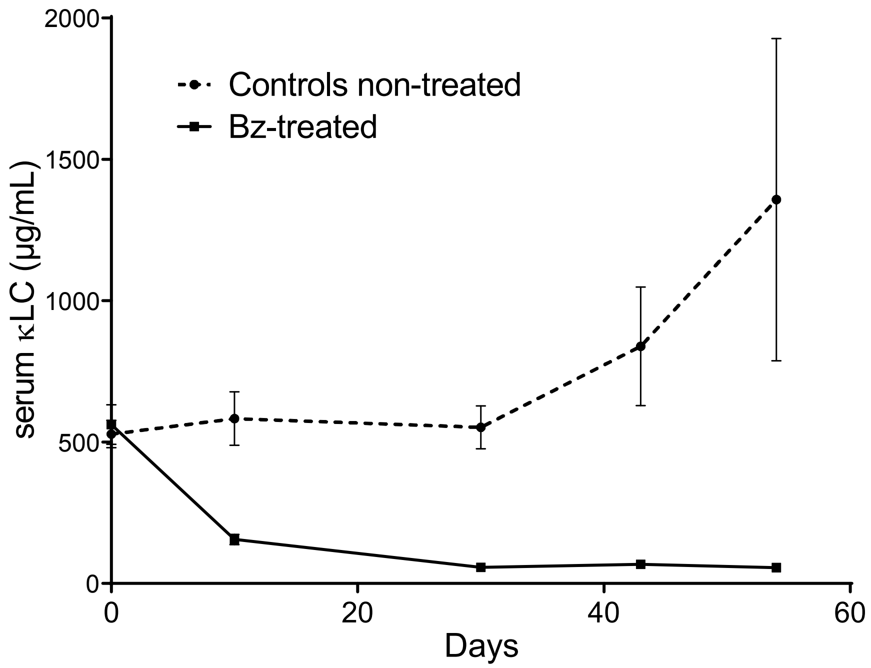


Supplemental Figure 8 : **Bortezomib/cyclophosphamide-based therapy efficacity in κF-DH mice.** Serum free light chains levels (in µ/mL) in treated and non-treated κF-DH mice during the course of the treatment.
